# FGF2 binds to the allosteric site (site 2) and activates αvβ3 integrin and FGF1 binds to site 2 but suppresses integrin activation by FGF2: a potential mechanism of anti-inflammatory action of FGF1

**DOI:** 10.1101/2024.04.17.589976

**Authors:** Yoko K Takada, Xueson Wu, David Wei, Samuel Hwang, Yoshikazu Takada

## Abstract

FGF1 is known as an anti-inflammatory and has suppresses insulin resistance. Its homologue FGF2 is pro-inflammatory. Mechanism of FGF1’s anti-inflammatory action and FGF2’s pro-inflammatory action are unknown. Several inflammatory cytokines (e.g., CX3CL1, CCL5, and CXCL12, and CD40L) bind to the classical ligand (RGD)-binding site (site 1) of integrin αvβ3. In addition, they bind to the allosteric site (site 2) of αvβ3, which is distinct from site 1, and allosterically activate αvβ3. Site 2 is involved in inflammatory signals since inflammatory lipid mediator 5-hydroxycholesterol binds to site 2 and induces integrin activation and inflammatory signals (e.g., TNF and IL-6 secretion). We thus hypothesized that FGF1 and FGF2 bind to site 2 and affect activation status of integrins. Here we describe that FGF2 bound to site 2 and allosterically activated αvβ3 integrin. Point mutations in the site 2-binding interface of FGF2 suppressed this activation, indicating that FGF2 binding to site 2 is required for inducing integrin activation. In contrast, FGF1 bound to site 2 but did not activate αvβ3, and instead suppressed integrin activation induced by FGF2, indicating that FGF1 acts as an antagonist of site 2. These findings suggest that FGF1’s anti-inflammatory action is mediated by blocking site 2. FGF1 has potential as an anti-inflammatory agent, but is not appropriate for long-term use since it is potent mitogen. A non-mitogenic FGF1 mutant (R50E), which is defective in binding to site 1 of αvβ3, suppressed αvβ3 activation by FGF2 as effectively as WT FGF1. We propose that FGF1 R50E has therapeutic potential for inflammatory diseases.

## Introduction

Integrins are a superfamily of αβ heterodimers and are receptors for extracellular matrix (ECM) (e.g., vitronectin, fibronectin, and collagen), cell surface proteins (e.g., VCAM-1, and ICAM-1), and soluble ligands (e.g., growth factors) (Takada et al. 2007). Antagonists to integrins have been shown to suppress FGF2-induced angiogenesis and tumor growth, indicating that integrins are required for FGF signaling (FGF-integrin crosstalk) (Brooks et al. 1994). We identified FGF1 and FGF2 as new ligands for integrin αvβ3 by virtual screening of protein data bank (PDB) using docking simulation with the integrin headpiece as a target (Mori et al. 2008, Mori et al. 2017). Docking simulation predicts that FGF1 and FGF2 bind to the classical ligand-binding site of αvβ3 (site 1) (Mori et al. 2008) (Mori et al. 2017). FGF1 binding to integrin αvβ3 induced integrin αvβ3-FGF1-FGFR1 ternary complex formation (Yamaji et al. 2010). The FGF1 mutant defective in integrin binding due to a point mutation in the predicted integrin-binding site (Arg50 to Glu, R50E) did not bind to integrin αvβ3 but still bound to FGFR and heparin and was defective in inducing sustained ERK1/2 activation, ternary complex formation, and inducing mitogenesis (Mori et al. 2008). The R50E FGF1 mutant was defective in mitogenesis and in inducing angiogenesis, and suppressed angiogenesis induced by WT FGF1 (dominant-negative effect) (Mori et al. 2013). We obtained similar result in FGF2 (Mori et al. 2017). The FGF2 mutants (K119E/R120E and K125E) in the integrin-binding interface of FGF2 were defective in signaling, ternary complex formation, and acted as dominant-negative antagonists (Mori et al. 2017). It has been reported that FGF2 and integrin α6β1 are important for maintaining the pluripotency of human pluripotent stem cells (hPSCs). It has recently been reported that integrin α6β1-FGF2-FGFR ternary complex formation is critical for maintaining the pluripotency of hPSCs (Cheng et al. 2024). The FGF2 K125E was incapable of inducing the hPSC properties, such as proliferation, ERK activity, and large focal adhesions at the edges of human induced pluripotent stem cells (hiPSCs) colonies. These findings indicate that FGF1 and FGF2 binding to integrin (site 1) and ternary complex formation is critical for their signaling functions.

It has been well established that integrin activation can be mediated by signals from inside the cells (inside-out signaling) (Shattil et al. 2010, Ginsberg 2014). We discovered, however, that several pro-inflammatory proteins such as CX3CL1 (fractalkine) (Fujita et al. 2014), CXCL12 (SDF-1) (Fujita et al. 2018, Takada et al. 2022), Rantes (CCL5) (Takada et al. 2022), and secreted phospholipase A2 type IIA (sPLA2-IIA) (Fujita et al. 2015), CD40L (Takada et al. 2021), and P-selectin (Takada et al. 2023) activated integrins independent of inside-out signaling. We found that this activation is induced by ligand binding to the allosteric ligand-binding site (site 2) of integrins. Site 2 is distinct from the classical ligand-binding site (site 1) and is on the opposite side of site 1 in the integrin headpiece (Fujita et al. 2014, Fujita et al. 2018, Takada et al. 2022). We showed that peptides from site 2 (site 2 peptides) bound to these allosteric activators and suppressed integrin activation, indicating that they are required to bind to site 2 for allosteric integrin activation. Since the allosteric activation is induced by inflammatory cytokines, there should be a link between allosteric integrin activation and inflammation. Recently, pro-inflammatory lipid mediator, 25-hydroxycholesterol, was found to bind to site 2 of integrins and activates integrins and induce inflammatory signals (e.g., secretion of IL-6 and TNF) (Pokharel et al. 2019), which verify the role of site 2 in allosteric integrin activation and inflammatory signals in inflammation.

FGF2 is pro-inflammatory and induces the expression of a wide repertoire of inflammation-related genes in endothelial cells, including pro-inflammatory cytokines/chemokines and their receptors, endothelial cell adhesion molecules, and components of the prostaglandin pathway (Presta et al. 2009). FGF2 expression is enhanced in endothelial precursor cells in deep vein thrombosis (Sun et al. 2020), suggesting that FGF2 is prothrombotic.

FGF1 is another member of the same subfamily as FGF2 (FGF1 subfamily) and is not stored in platelet granules. Previous studies showed that FGF1 is anti-inflammatory (Fan et al. 2019); however, the mechanism of anti-inflammatory effects of FGF1 is unclear. Also, FGF1 is shown to lower blood glucose levels in diabetic mice but the mechanism of this action is unknown (Suh et al. 2014). FGF1 significantly prevented the development of nonalcoholic fatty liver disease (NAFLD) and diabetic nephropathy (DN) (Liu et al. 2016, Liang et al. 2018). Also, FGF1 is shown to lower blood glucose levels in diabetic mice but the mechanism of this action is unknown (Suh et al. 2014).

In the present study, we studied if FGF2 (pro-inflammatory) and FGF1 bind to site 2 of αvβ3 and affect the activation status of this integrin. We describe here that FGF2 activated αvβ3 by binding to site 2. FGF1 also bound to site 2 but did not activate αvβ3, and instead suppressed integrin activation induced by FGF2, indicating that FGF1 act as an antagonist of site 2. We propose a model, in which pro-inflammatory action of FGF2 is mediated by binding to site 2, and anti-inflammatory action of FGF1 is mediated by inhibiting site 2-mediated αvβ3 activation by FGF2. Non-mitogenic FGF1 R50E also suppressed activation of αvβ3 by FGF2 in cell-free conditions. We propose that site 2 is a novel therapeutic target for inflammatory diseases.

## Materials and Methods

### Materials

The truncated fibrinogen γ-chain C-terminal domain (γC399tr) was generated as previously described (Yokoyama et al. 1999). The protein was synthesized in E. coli BL21 and purified using glutathione affinity chromatography. The protein was synthe-sized in E. coli BL21 and purified using glutathione affinity chromatography.

FGF1 (Mori et al. 2008) and FGF2 (Mori et al. 2017) were synthesized as previously described.

Cyclic β3 site 2 peptide fused to GST-The 29-mer cyclic β3 site 2 peptide C260-RLAGIV[QPNDGSHVGSDNHYSASTTM]C288 (C273 is changed to S) was synthesized by inserting oligonucleotides encoding this sequence into the BamHI-EcoRI site of pGEX-2T vector. The positions of Cys residues for disulfide linkage were selected by using Disulfide by Design-2 (DbD2) software (http://cptweb.cpt.wayne.edu/DbD2/) (Craig and Dombkowski 2013). It predicted that mutating Gly260 and Asp288 to Cys disulfide-linked cyclic site 2 peptide of β3 does not affect the conformation of the original site 2 peptide sequence QPNDGSHVGSDNHYSASTTM in the 3D structure. We found that the cyclic site 2 peptide bound to CX3CL1 and sPLA2-IIA to a similar extent to non-cyclized β3 site 2 peptides in ELISA-type assays (data not shown). We designed the corresponding cyclic β1 peptide (C268-KLGGIVLPNDGQSHLENNMYTMSHYYC295, 28-mer cyclic β1 peptide) in which C281 is converted to S. We synthesized the proteins in BL21 cells and purified using glutathione-Sepharose affinity chromatography.

### Activation of soluble αvβ3 by FGF2

ELISA-type binding assays were performed as described previously (Fujita et al. 2018). Briefly, wells of 96-well Immulon 2 microtiter plates (Dynatech Laboratories, Chantilly, VA) were coated with 100 μl 0.1 M PBS containing γC399tr for αvβ3 for 2 h at 37°C. Remaining protein-binding sites were blocked by incubating with PBS/0.1% BSA for 30 min at room temperature. After washing with PBS, soluble recombinant αIbβ3 or αvβ3 (1 μg/ml) in the presence or absence of FGF1 or FGF2 was added to the wells and incubated in Hepes-Tyrodes buffer (10 mM HEPES, 150 mM NaCl, 12 mM NaHCO_3_, 0.4 mM NaH_2_PO_4_, 2.5 mM KCl, 0.1% glucose, 0.1% BSA) with 1 mM CaCl_2_ for 1 h at room temperature. After unbound αvβ3 was removed by rinsing the wells with binding buffer, bound αvβ3 was measured using anti-integrin β3 mAb (AV-10) followed by HRP-conjugated goat anti-mouse IgG and peroxidase substrates. For the time-course experiments, WT FGF2 and soluble αvβ3 was incubated for 1–60 min instead of 1 h.

### Binding of site 2 peptide to FGF1 and FGF2

ELISA-type binding assays were performed as described previously (Fujita et al. 2014). Briefly, wells of 96-well Immulon 2 microtiter plates (Dynatech Laboratories, Chantilly, VA) were coated with FGF1 or FGF2 (10 μg/ml) in 100 μl in PBS for 2 h at 37°C. Remaining protein-binding sites were blocked by incubating with PBS/0.1% BSA for 30 min at room temperature. After washing with PBS, GST-β3 peptide (100 μg/ml) was added to the wells and incubated in PBS/0.5% Tween 20 for 1 h at room temperature. After unbound site 2 peptide was removed by rinsing the wells with PBS/0.5% Tween 20, bound site 2 peptide was measured using HRP-conjugated anti-GST antibody.

### Docking simulation

Docking simulation of interaction between FGF2 and integrin αvβ3 (closed headpiece form, PDB code 1JV2) was performed using AutoDock3 as described previously (Ieguchi et al. 2010). We used the headpiece (residues 1–438 of αv and residues 55– 432 of β3) of αvβ3 (closed form, 1JV2.pdb). Cations were not present in integrins during docking simulation, as in the previous studies using αvβ3 (open headpiece form, 1L5G.pdb) (Mori et al. 2008, Saegusa et al. 2008). The ligand is presently compiled to a maximum size of 1024 atoms. Atomic solvation parameters and fractional volumes were assigned to the protein atoms by using the AddSol utility, and grid maps were calculated by using AutoGrid utility in AutoDock 3.05. A grid map with 127 × 127 × 127 points and a grid point spacing of 0.603 Å included the headpiece of αvβ3. Kollman ‘united-atom’ charges were used. AutoDock 3.05 uses a Lamarckian genetic algorithm (LGA) that couples a typical Darwinian genetic algorithm for global searching with the Solis and Wets algorithm for local searching. The LGA parameters were defined as follows: the initial population of random individuals had a size of 50 individuals; each docking was terminated with a maximum number of 1 × 10^6^ energy evaluations or a maximum number of 27 000 generations, whichever came first; mutation and cross-over rates were set at 0.02 and 0.80, respectively. An elitism value of 1 was applied, which ensured that the top-ranked individual in the population always survived into the next generation. A maximum of 300 iterations per local search were used. The probability of performing a local search on an individual was 0.06, whereas the maximum number of consecutive successes or failures before doubling or halving the search step size was 4.

### Other methods

Treatment differences were tested using ANOVA and a Tukey multiple comparison test to control the global type I error using Prism 10 (Graphpad Software).

## Results and Discussion

Previous studies identified cytokines (e.g., fractalkine, CCL5, CXCL12, CD40L, sPLA2-IIA, and FGF2) bind to the classical ligand (RGD)-binding site (site 1) by virtual screening of protein data bank using the headpiece of αvβ3 (1L5G.pdb, open headpiece form) as a target (See introduction). We also found that several pro-inflammatory cytokines bind to the allosteric site (site 2), which is distinct from site 1, and allosterically activated integrin αvβ3 in 1 mM Ca^2+^ (See introduction). We showed that FGF1 (anti-inflammatory) and FGF2 (pro-inflammatory) bind to site 1 (Mori et al. 2008, Mori et al. 2017). It has been well established that FGF1 is anti-inflammatory, but the mechanism of its anti-inflammatory action is unknown. it is unclear if FGF1 and FGF2 bind to site 2 and activate integrin αvβ3. We hypothesized that FGF1 and FGF2 may interact with site 2 in a different manner, leading to anti- or pro-inflammatory actions.

### FGF2 binds to site 2 and activates integrin αvβ3 in an allosteric manner

The docking simulation of interaction between FGF2 (1BFG.pdf) and αvβ3 (closed headpiece form, 1JV2.pdb) predicts that FGF2 binds to site 2 (docking energy -20.1 Kcal/mol) (Fig. 1a). The simulation predicted that FGF2 activates αvβ3 in an allosteric manner. Amino acid residues that are involved in FGF2-site 2 interaction is shown in Table 1. We studied if FGF2 activated integrin αvβ3 in 1 mM Ca^2+^ in cell-free conditions in ELISA-type integrin activation assays. Wells of 96-well microtiter plate were coated with a fibrinogen fragment (γC399), a specific ligand for αvβ3, and incubated with soluble integrin αvβ3 in 1 mM Ca^2+^ in the presence of FGF2. We found that FGF2 enhanced the binding of soluble αvβ3 to immobilized γC399 in a dose-dependent manner (Fig. 1b).

**Table 1.** Amino acid residues of FGF2 involved in binding to site 2 of αvβ3 (1JV2.pbd) predicted by docking simulation.

| FGF2 (1BFG) -20.1 | $\alpha\text{v}$ (1JV2) | $\beta 3$ |
| --- | --- | --- |
| Asp-19, **Arg33**, His35, Pro36, Asp37, Gly38, **Arg39**, Val43, Glu45, **Lys46**, Ser47, Asp48, Pro49, His50, Gln56, **Lys66**, Val68, Ser69, Ala70, Asn71, **Arg72**, Tyr73, **Lys77**, **Arg81**, Leu83, Ala84, Ser85, **Lys86**, Ser87, Val88, Thr89, Asp89, Asp90, Phe93, **Lys110**, | Glu15, Asn44, Gly49, Ile50, Val51, Glu52, Asn77, Asp83, Phe88, Ser90, His91, Arg122, | Pro160, Val161, Ser162, Met165, Ile167, Ser168, Glu171, Glu174, Asn175, Pro186, Met187, Lys235, <u>Val275, Gly276, Ser277, Asp278, His280, Tyr281, Ser282, Ala283, Thr285, Thr286</u> |

Amino acid residues within 0.6 nM between FGF2 and $\alpha\beta 3$ were selected using PDB viewer (version 4.1) (Swiss Institute of Bioinformatics, Basel, Swiss). Amino acid residues in $\beta 3$ that are in the cyclic site 2 peptide are underlined.

**Fig. 1.**
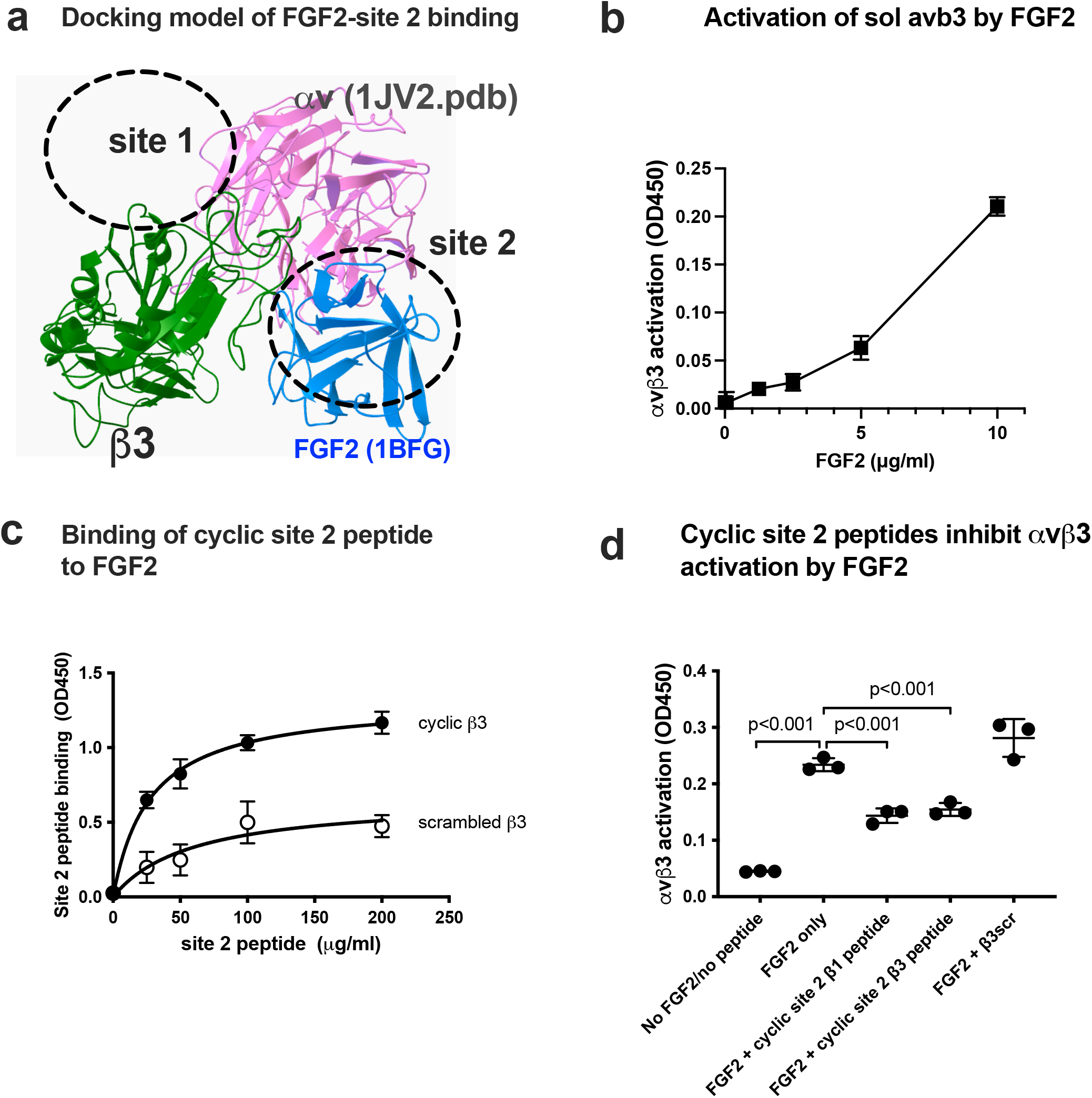
FGF2 binds to site 2 of integrin and αvβ3 and activates integrins (act as an agonist). (a) Docking simulation of interaction between FGF2 (1BFG.pdb) and αvβ3 (1JV2.pdb). FGF2 was predicted to bind to site 2. (b) FGF2 activates soluble αvβ3 in cell-free conditions. The fibrinogen fragment (γC399tr), a specific ligand to αvβ3, was immobilized to wells of 96-well microtiter plate and incubated with soluble αvβ3 (1 μg /ml) in 1 mM Ca^2+^ in the presence of FGF2, and bound αvβ3 was measured. The data shows that FGF2 activated αvβ3. (c) Cyclic site 2 peptides from β3 binds to FGF2 better than control β3 scrambled peptide. ELISA-type binding assay was performed as described in the methods section. (d) Cyclic site 2 peptides inhibit FGF2-induced αvβ3 activation. Bound integrin was measured. ANOVA using Prism 10 was used for statical analysis (n=3).

Previous studies showed that several integrin ligands that bind to site 2 and activates integrins bound to peptides from site 2 (of β1 and β3) (Fujita et al. 2014, Fujita et al. 2015, Fujita et al. 2018, Takada et al. 2021, Takada et al. 2022, Yoko K Takada 2022). We found that cyclic peptides derived from site 2 of β3 bound to FGF2 at higher levels than control scrambled β3 peptide, indicating that FGF2 is required to bind to site 2 to activate αvβ3 (Fig. 1c). We found that cyclic site 2 peptides from integrin β1 or β3 suppressed αvβ3 activation by FGF2 (Fig 1d), indicating that activation of αvβ3 by FGF2 requires FGF2 binding to site 2.

### Point mutations of the predicted site 2-binding interface in FGF2 block FGF2-mediated activation of αvβ3

To further show how FGF2 binds to site 2 of αvβ3, we generated FGF2 mutants defective in site 2 binding. The docking simulation of interaction between FGF2 (1BFG.pdf) and αvβ3 (1JV2.pdb) predicts that amino acid residues Lys66, Arg72, Lys77, Lys86, and Lys110 of β3 interact with αvβ3 (docking energy -20.5 Kcal/mol) (Fig. 3a). The docking simulation predicts that amino acids Lys66, Arg72, Lys77, Lys86, and Lys110 of β3 are in the site 2-binding interface of FGF2 (Fig. 2a). We found that mutation of these amino acids to Glu (K66E, K72E, K77E, K86E, and K110E) effectively suppressed integrin activation by FGF2 (Fig. 2b), indicating that the docking model is correct and FGF2 binding to site 2 is required for activation of αvβ3. These mutations did not inhibit FGF2 binding to site 1 of αvβ3 in 1 mM Mn^2+^ (Fig. 2c), which is consistent with the previous findings that these mutations are not in the site 1-binding interface of FGF2 (Mori et al. 2017). We previously showed that FGF2 binds to the classical ligand-binding site (site 1) and the FGF2 mutants (K119E/R120E and K125E) in the integrin-binding interface of FGF2 were defective in binding to αvβ3, in signaling, ternary complex formation, and acted as dominant-negative antagonists (Mori et al. 2017). These mutations are predicted to be not in the site 2-binding interface (Fig. 2a)(Table 1). We studied if FGF2 mutants defective in site 1 binding activate αvβ3. The K119E/R120E are K125E mutants in the site 1-binding site of FGF2 induced αvβ3 activation by FGF2 (Fig. 2d), indicating that site 2 and site 1-binding interfaces in FGF2 are distinct (Fig. 2d). These findings suggest that FGF2 mutants defective in site 2 binding are defective in integrin activation, but not in site 1 binding. Likewise, FGF2 mutants defective in site 1 binding still bind to site 2 and activate αvβ3.

**Fig. 2.**
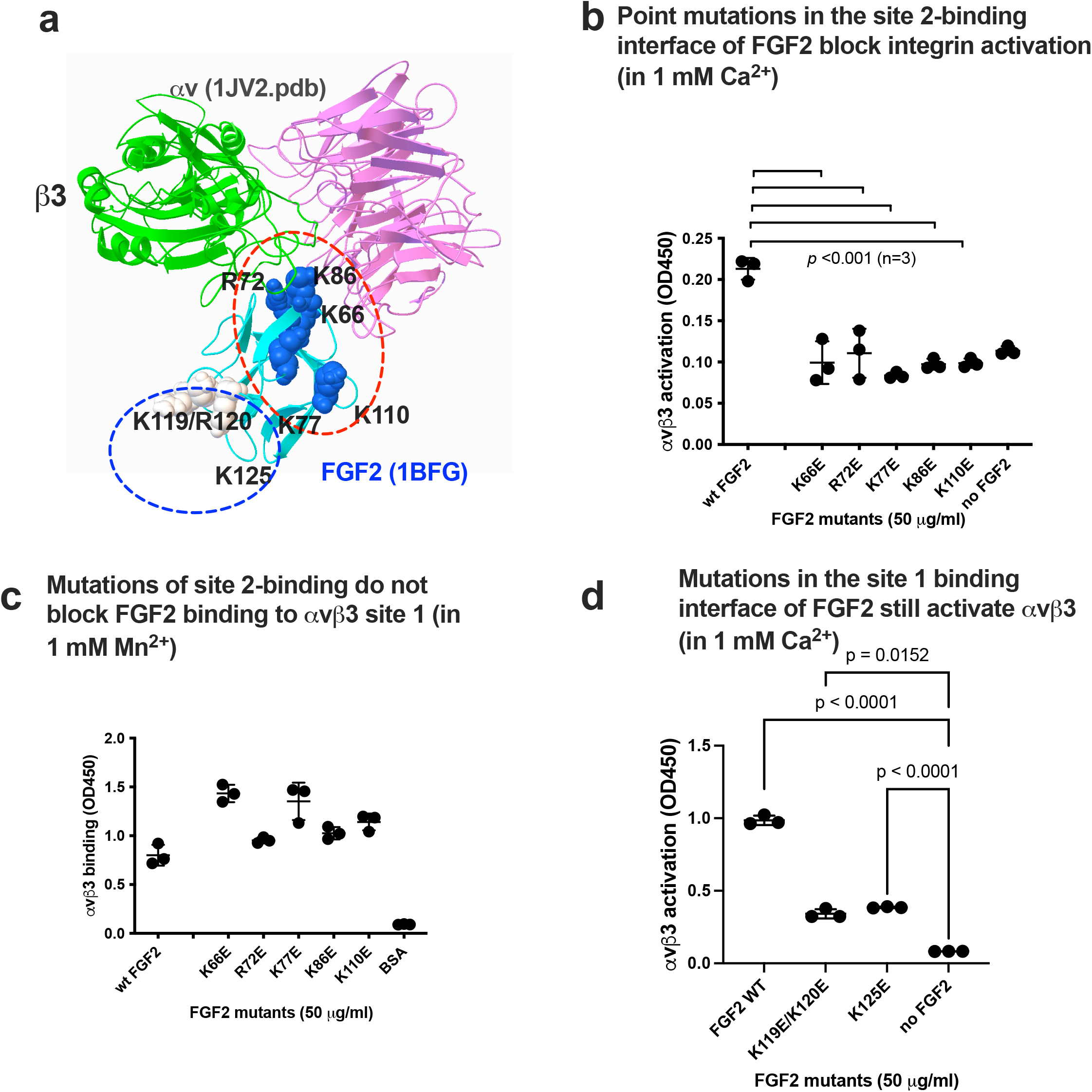
Point mutations in site 2-binding interface of FGF2 effectively reduce activation of integrin αvβ3 by FGF2. (a). Positions of amino acid residues involved in site 1 and site 2 binding predicted by docking simulations. K119E/R120E and K125E mutations suppressed FGF2 binding to integrin site 1 of αvβ3 and thereby suppressed FGF2 mitogenicity (Mori et al. 2017), but still bind to site 2. (b) FGF2 with point mutations in the predicted site 2-binding interface of FGF2 did not activate integrin αvβ3. Activation assays were performed as described in Fig.1. (c) Point mutations in the predicted site 2-binding interface did not affect FGF2 binding to site 1 in 1 mM Mn^2+^. Wells of 96-well microtiter plate were coated with FGF2 WT and mutants and incubated with soluble αvβ3 in 1 mM Mn^2+^. Bound αvβ3 was quantified using anti-β3 and anti-mouse IgG conjugated with HRP. (d) Mutations in the site 1-binding interface of FGF2 still activate integrin αvβ3. Activation assays were performed as described in Fig.1. The data is shown as means +/- SD in triplicate experiments.

**Fig. 3.**
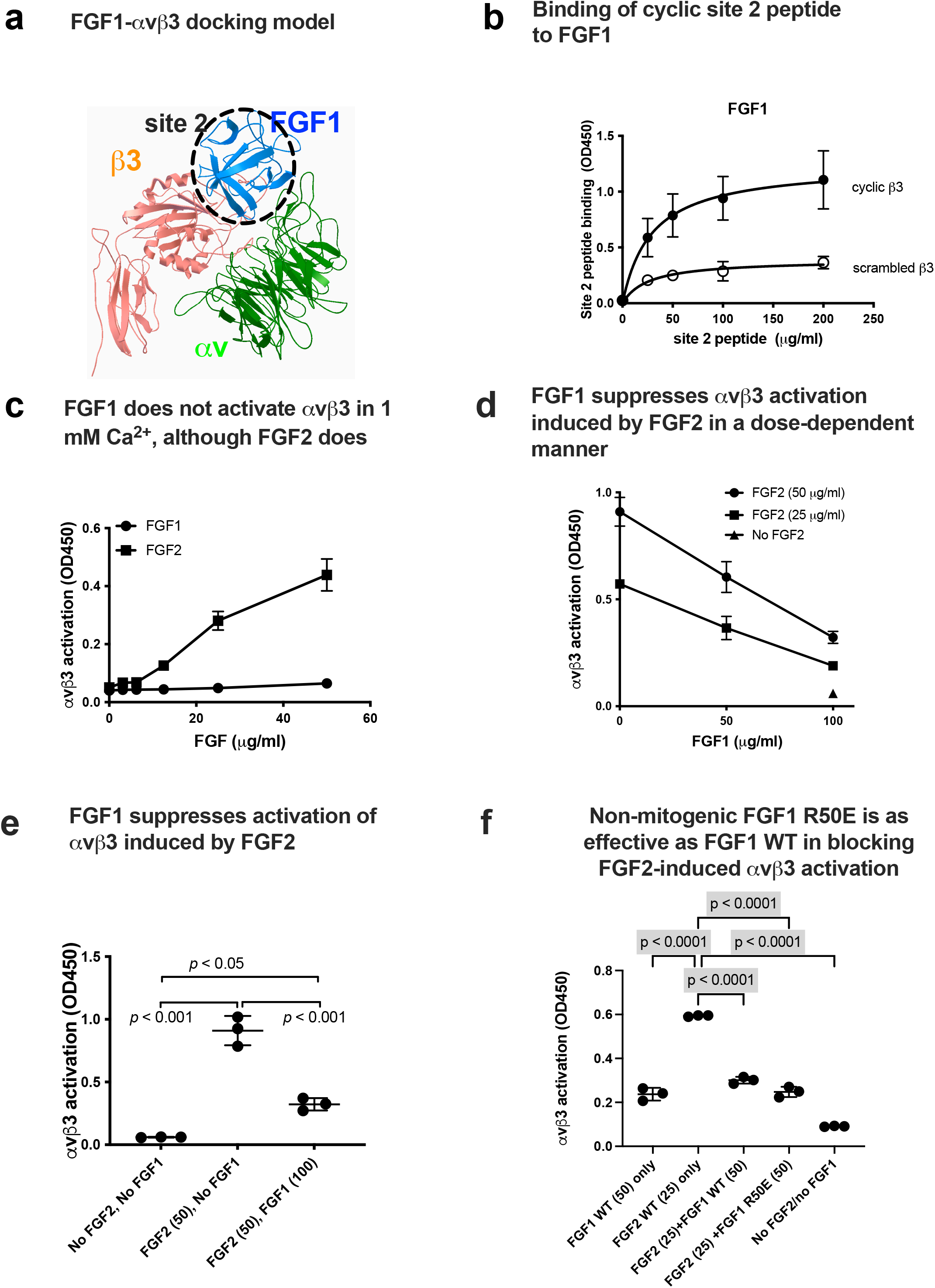
FGF1 binds to site 2 but did not activate αvβ3. FGF1 suppressed integrin activation by FGF2. (a) Docking simulation of interaction between FGF1 (AXM1.pdb) and αvβ3 (1JV2.pdb). FGF1 is predicted to bind to site 2. Docking simulation was performed in the methods section. Amino acid residues that are predicted to be involved in the interaction were described in Table 2. (b) Cyclic site 2 peptides from β3 binds to FGF1 better than control β3 scrambled peptide. ELISA-type binding assay was performed as described in the methods section. (c) FGF2 allosterically activated soluble αvβ3 in a dose-dependent manner, but FGF1 did not. Wells of 96-well microtiter plate were coated with γC399tr, a specific ligand for αvβ3, and incubated with soluble αvβ3 in 1 mM Ca^2+^. Bound αvβ3 was quantified using anti-β3. (e).(d) FGF1 inhibits integrin activation by FGF2. Soluble αvβ3 (1 μg /ml) was incubated with immobilized ligand (γC399tr for αvβ3) in the presence of FGF2 and/or FGF1 in 1 mM Ca^2+^. (f) Non-mitogenic FGF1 mutant (R50E) suppressed integrin activation by FGF2 at a level comparable to that of WT FGF1 in Ca^2+^.

### FGF1 binds to site 2 but does not activate integrin αvβ3 and instead FGF1 suppresses αvβ3 activation by FGF2

Docking simulation of interaction between FGF1 and closed headpiece form of αvβ3 (1JV2.pdb) (Docking energy -22.4 Kcal/mol) predicts that FGF1 binds to site 2 (Fig. 3a). Consistently, we found that cyclic site 2 peptide from integrin β3 bound to FGF1 better than control scrambled β3 peptide (Fig. 3b), suggesting that FGF1 binds to site 2. So, it is predicted that FGF1 allosterically activates αvβ3. Unexpectedly, we found that FGF1 did not activate αvβ3 under the conditions in which FGF2 activated αvβ3 (Fig. 3c).

**Table 2.**
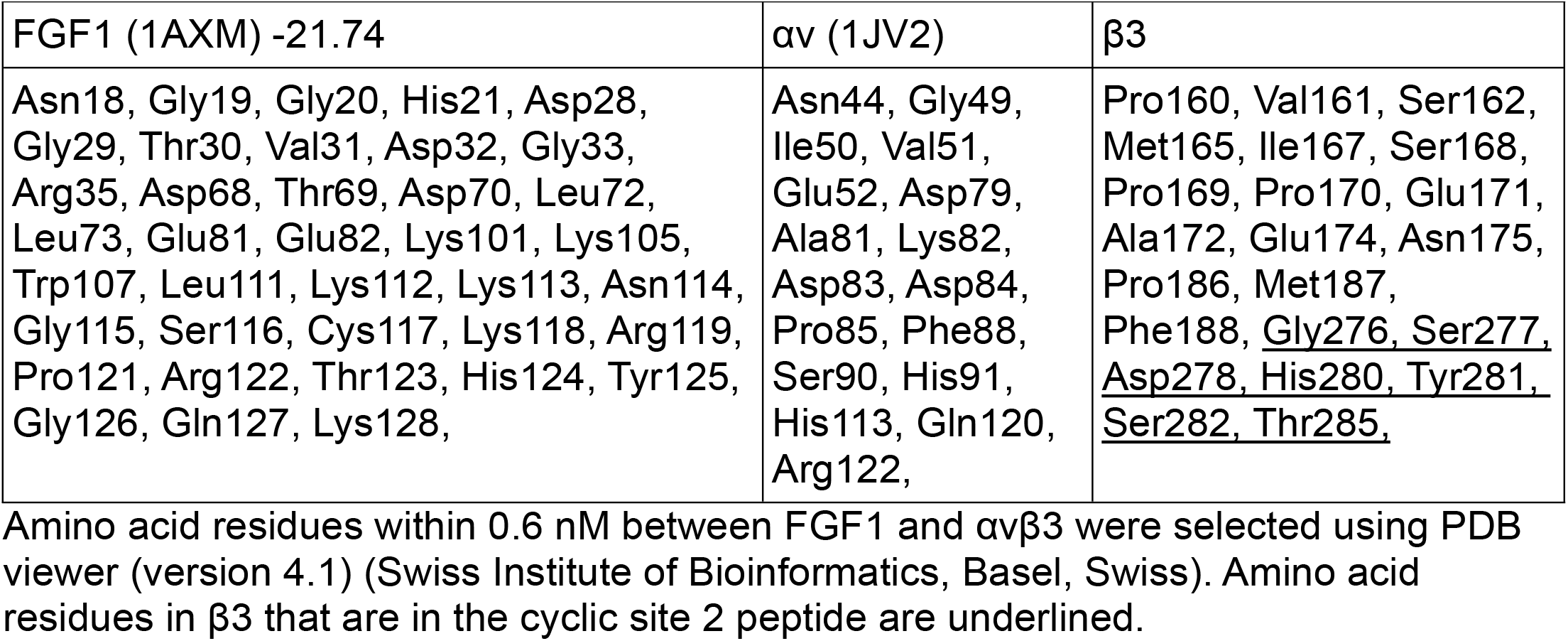
Amino acid residues of FGF1 involved in binding to site 2 of αvβ3 (1JV2.pbd) predicted by docking simulation.

| FGF1 (1AXM) -21.74 | $\alpha\text{v}$ (1JV2) | $\beta 3$ |
| --- | --- | --- |
| Asn18, Gly19, Gly20, His21, Asp28, Gly29, Thr30, Val31, Asp32, Gly33, Arg35, Asp68, Thr69, Asp70, Leu72, Leu73, Glu81, Glu82, Lys101, Lys105, Trp107, Leu111, Lys112, Lys113, Asn114, Gly115, Ser116, Cys117, Lys118, Arg119, Pro121, Arg122, Thr123, His124, Tyr125, Gly126, Gln127, Lys128, | Asn44, Gly49, Ile50, Val51, Glu52, Asp79, Ala81, Lys82, Asp83, Asp84, Pro85, Phe88, Ser90, His91, His113, Gln120, Arg122, | Pro160, Val161, Ser162, Met165, Ile167, Ser168, Pro169, Pro170, Glu171, Ala172, Glu174, Asn175, Pro186, Met187, Phe188, <u>Gly276, Ser277, Asp278, His280, Tyr281, Ser282, Thr285,</u> |

Amino acid residues within 0.6 nM between FGF1 and $\alpha\beta 3$ were selected using PDB viewer (version 4.1) (Swiss Institute of Bioinformatics, Basel, Swiss). Amino acid residues in $\beta 3$ that are in the cyclic site 2 peptide are underlined.

Notably, we found that excess (2x) FGF1 suppressed αvβ3 activation by FGF2 (Figs. 3d and 3e) in a dose-dependent manner, indicating that FGF1 binds to site 2 and acts as an inhibitor of allosteric activation by FGF2 (site 2 antagonist). FGF1’s antagonistic action for site 2-mediated integrin activation is a potential mechanism of its anti-inflammatory action.

### Non-mitogenic FGF1 R50E mutant suppresses αvβ3 activation by FGF2 at a comparable level of WT FGF1

Since WT FGF1 is a potent mitogen and, non-mitogenic FGF1 has been sought. We showed that FGF1 R50E (defective in site 1 binding) is defective in inducing integrin-FGF1-FGFR1 ternary complex, and defective in inducing sustained ERK1/2 activation and AKT activation (Mori et al. 2008). R50E binds to FGFR and heparin, but not to site 1 of integrins. FGF R50E suppressed tumorigenesis and angiogenesis in vivo (Mori et al. 2013). In the present study, we found that non-mitogenic FGF1 R50E inhibited activation of αvβ3 by FGF2 in solution as effectively as WT FGF1 (Fig. 3f). This indicates that FGF1 R50E has potential as a therapeutic.

### Site 2 as a potential target for anti-inflammatory agents

The present study showed that FGF1 and FGF2 also bind to site 2 of αvβ3. FGF2 bound to site 2 and activated αvβ3, as predicted, indicating that FGF2 acts as an agonist of site 2 (Fig. 4). However, FGF1 bound to site 2 of αvβ3, but did not activate αvβ3. Instead, FGF1 suppressed activation of αvβ3 induced by FGF2, indicating that FGF1 acts as an antagonist of site 2. It has been well established that FGF1 is anti-inflammatory, but the mechanism of anti-inflammatory action of FGF1 is unknown. Previous studies showed that FGF1 remarkably lowered levels of several serum inflammatory cytokines and impeded the inflammatory response (Suh et al. 2014, Liang et al. 2018). We propose that FGF1’s anti-inflammatory actions may be mediated by blocking inflammatory signals through site 2.

**Fig. 4.**
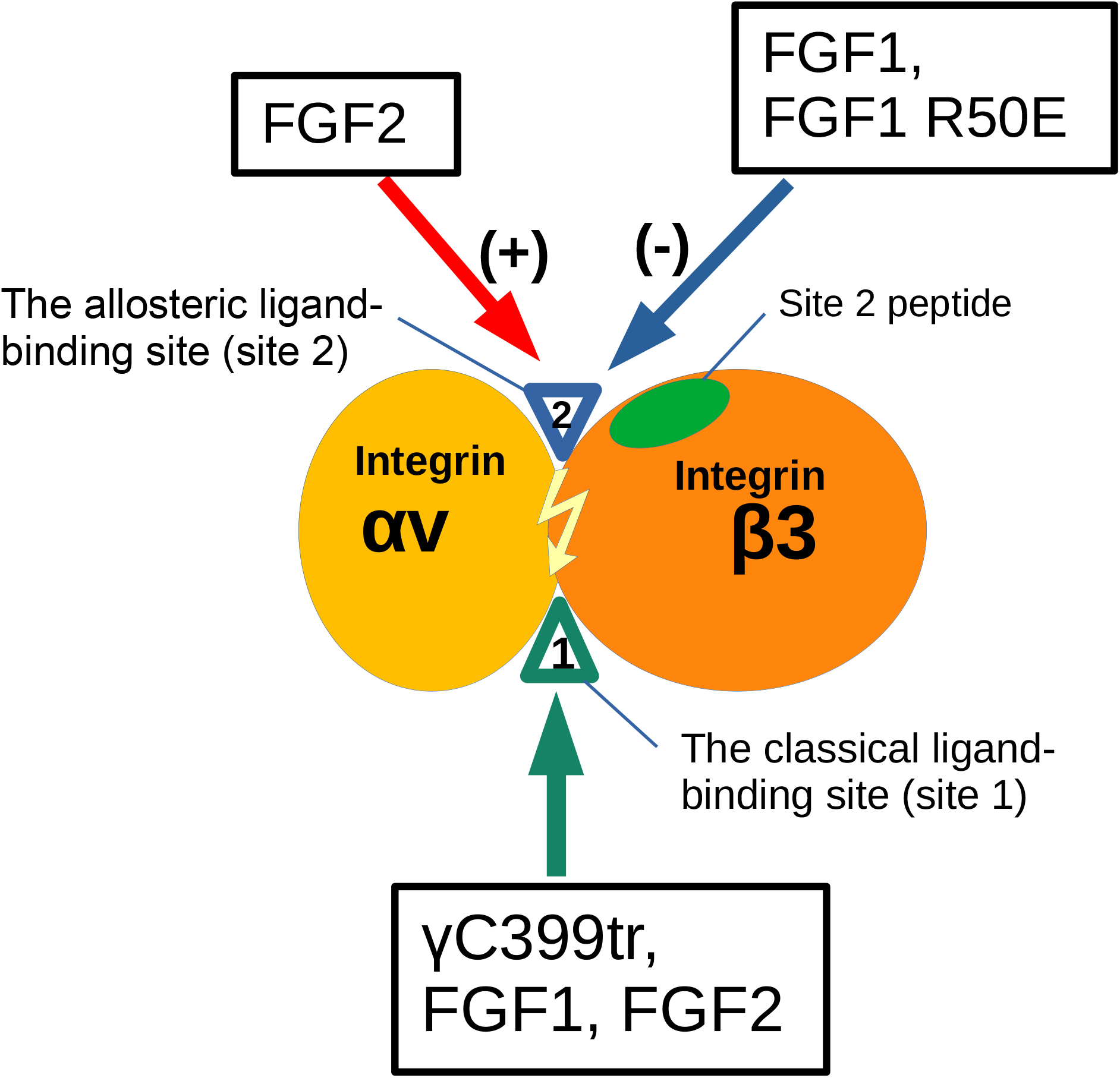
Agonistic action of FGF2 and antagonistic action of FGF1 to the allosteric site (site 2) of β3 integrins. (b) Fibrinogen γ-chain fragment (γC399tr) binds specifically to integrin αvβ3. We previously showed that FGF1 and FGF2 bind to site 1 of αvβ3 (Mori et al. 2008, Mori et al. 2017). FGF2 binds to site 2 and induces integrin activation. FGF1 also binds to site 2 and blocks FGF2-induced integrin activation. This is a potential mechanism of anti-inflammatory action of FGF1. Non-mitogenic FGF1 R50E also inhibits FGF2-induced integrin activation, and has potential as an anti-inflammatory agent.

Since WT FGF1 is a potent mitogen and, non-mitogenic FGF1 has been sought. Deletion of N-terminal residues of FGF1 (amino acid residues 21-27), which lacks nuclear translocation signal, has been shown to be non-mitogenic and still has anti-inflammatory activity (Imamura et al. 1990). It has been reported that this N-terminal truncated FGF1 induced angiogenesis (Zou et al. 2020). FGF1 R50E has been well characterized as non-mitogenic FGF1. R50E binds to FGFR and heparin, but not to site 1 of integrins. R50E is defective in inducing integrin-FGF1-FGFR1 ternary complex, and defective in inducing sustained ERK1/2 activation and AKT activation (Mori et al. 2008). We showed that non-mitogenic FGF1 R50E (defective in site 1 binding) suppressed tumorigenesis and angiogenesis in vivo (Mori et al. 2013). This indicates that FGF1 R50E has potential as a therapeutic for inflammatory and thrombotic diseases, and insulin resistance.

